# Prevalence and Diversity of Haemosporidian-Associated Matryoshka RNA Viruses in a Natural Population of Wild Birds

**DOI:** 10.1101/2024.10.23.618521

**Authors:** Carlos W. Esperanza, Caroline E. Faircloth, Scott W. Roy, Ravinder N.M. Sehgal

## Abstract

Matryoshka RNA viruses (MaRNAV) have previously been detected using bioinformatics and limited PCR approaches. They are associated with haemosporidian parasite infections, yet their prevalence and diversity in wild bird populations remains largely unknown. To investigate the prevalence of MaRNAV, we examined blood samples collected from wild passerine birds and raptors in the San Francisco Bay Area. Samples were first screened for haemosporidian infections followed by RNA sequencing (RNAseq) and reverse transcriptase (RT) PCR to detect MaRNAV. Our analyses identified two novel MaRNAV (MaRNAV-5 and −6) infecting various bird species harboring diverse *Haemoproteus* and *Leucocytozoon* species and lineages. MaRNAV-5, associated with *Haemoproteus*, exhibited 71.3% amino acid identity to MaRNAV-4, also associated with *Haemoproteus*, and was found across 15 different passerine species. MaRNAV-6, linked to *Leucocytozoon*, shared 72.9% identity with *Leucocytozoon-*associated MaRNAV-3 and was exclusively found in 4 raptor species. Prevalence was 44.79% for MaRNAV-5 in haemosporidian-infected passerines and 22.22% for MaRNAV-6 in haemosporidian-infected raptors. These viruses were not found in uninfected birds similarly tested via RNAseq and RT-PCR and were consistently only found in birds infected with haemosporidia. Sanger sequencing revealed high similarity of viral sequences across different bird species, even of different orders. Our findings indicate a high prevalence of MaRNAV among local wild birds, as well as their potential specificity to haemosporidia genera, suggesting potential impacts on their health and ecology. We propose a potential life-cycle model for this group of viruses where the insect vector is the primary host, and the haemosporidian parasite acts as the virus’ “vector” to reach its next host. Further research is needed to determine the impact of these viruses in avian systems.

## 1. Introduction

Haemosporidian parasites (Apicomplexa) are a diverse group of single-celled protozoa that invade the red blood cells of mammals, reptiles, and avian hosts (Garnham, 1966; Telford Jr., 2009; Valkiūnas, 2005). These parasites have been extensively studied and are found in most regions of the world (Clark et al., 2014). Avian-specific haemosporidian parasites consist of many species in the genera *Plasmodium*, *Haemoproteus*, and *Leucocytozoon*, which have distinct vectors, life cycles, and disease manifestations (Valkiūnas, 2005). Despite these differences, acute parasitemia in an avian host leads to avian malaria, and similar malaria-type infections, which have had some historically devastating impacts on wild bird populations with no natural exposure or immunity to avian malaria (Atkinson et al., 1995; Valkiūnas, 2005). Typically, if individuals survive acute infection, they will likely live with less severe chronic infections for the rest of their lives, with varying amounts of subsequent parasite recurrence (Valkiūnas, 2005; Himmel et al., 2024). However, some species of haemosporidia show variation in infectivity, with parasitemia being barely detectable in some bird species, and higher risk of major acute infection being linked to naïve bird communities (Palinauskas et al., 2008; Dimitrov et al., 2015). Recent genomic research on avian haemosporidia has seen major advances, including the complete genomic sequencing of the *Plasmodium relictum*, and several analyses of the effects that parasitic infection has on hosts’ transcriptomes by means of experimental inoculation (Ellis et al., 2022; Paxton et al., 2023; Videvall et al., 2020, 2021). Recently, with the advancement of high throughput sequencing and bioinformatics, several viruses have been described that appear to be associated with these parasites. Despite these technological advances, little is known regarding the biology of these viruses (Charon et al., 2019; Rodrigues et al, 2022).

Viruses are extremely abundant and can infect all types of life (Forterre, 2010). Of the known viruses, RNA viruses make up most of all recognized viral species, with new viruses being described yearly (Woolhouse et al., 2013). Among the viruses with the simplest genomes are the narnaviruses (Narnaviridae) (Dinan et al., 2020). Narnavirus genomes consist of a single-segmented positive-sense ssRNA, typically in the range of 2.3 to 3.6 kilobases (kb), that encodes an RNA-dependent RNA polymerase (RdRp) (Dinan et al., 2020). While most narnavirus genomes are single-segmented, recent findings have described more complex narnaviruses with additional putative segments, such as Culex narnavirus 1 (CxNV1) and Zhejiang mosquito virus 3, both originating in protist hosts (Retallack et al., 2021). Some narnavirus genomes have the unique feature of being ambigrammatic, with the reverse complement of their genomic sequences coding large open reading frames (ORFs) (DeRisi et al., 2019; Retallack et al., 2021). Viruses that specifically infect parasitic protozoa are known as Parasitic Protozoan Viruses (PPVs) and are all single-stranded (ss) or double-stranded (ds) RNA viruses (Wang & Wang, 1991). To date, PPVs have been characterized in *Trichomonas vaginalis*, *Giardia lamblia, Cryptosporidium parvum*, *Leishmania* spp., *Blechomonas* spp., and more recently, *Toxoplasmosis gondii* (Grybchuk et al., 2018; Gupta et al., 2024; Khramstov et al., 1997; Wang et al., 1986; Widmer et al., 1989). The presence of PPVs can significantly alter aspects of the parasite’s pathogenicity by affecting the animal host’s immune response (Gómez-Arreaza et al., 2017; Heeren et al., 2023; Zhao et al., 2023, Gupta et al., 2024). Leishmania RNA virus 1 (LRV1), Cryptosporidium parvum virus 1 (CSpV1), and Trichomonas vaginalis virus (TVV) have all been shown to weaken the host’s defenses against their respective parasite by triggering a type I interferon (IFN) inflammatory response in their hosts (Ives et al., 2011; Fichorova et al., 2012; de Carvalho et al., 2019; Rada et al., 2022; Deng et al., 2023). On the other hand, some evidence shows that Giardia lamblia virus 1 (GlV1) limits the growth of *G. lamblia* in its host, thereby mitigating infection (Miller et al., 1988).

In 2019, the first PPVs associated with haemosporidian parasites were described (Charon et al., 2019). Taking a meta-transcriptomic approach, they tested blood samples collected from human patients in Eastern Malaysia infected with various species of *Plasmodium*: *P. falciparum*, *P. vivax*, and *P. knowlesi*, and exhibiting clinical symptoms of malaria (Charon et al., 2019). This led to the identification of a novel viral sequences encoding an RdRp and a hypothetical protein with no known function that was restricted to the *P. vivax* samples, suggesting that the virus was species-specific (Charon et al., 2019). Further analysis of additional meta-transcriptomes from geographically diverse areas available on the NCBI SRA detected viral sequences that mapped to this RdRp and second sequence, all restricted to *P. vivax*-infected samples. This novel virus was named Matryoshka RNA virus 1 (MaRNAV-1) because of its Russian doll-like nature of a virus infecting a parasite infecting a host (Charon et al., 2019). Expanding their research to 12 *Leucocytozoon*-infected Australian avian meta-transcriptomes, and using MaRNAV-1 sequences as a reference, they described a second Matryoshka virus, MaRNAV-2, detected in 8 samples infected with *Leucocytozoon* (Charon et al., 2019). In 2021, further metatranscriptomic research has since identified two additional MaRNAV: MaRNAV-3, associated with *Leucocytozoon*, and MaRNAV-4, associated with *Haemoproteus* (Rodrigues et al., 2022). Phylogenetic analysis has revealed that the RdRps of these viruses are closely related to the RdRp of narnaviruses, with a major differentiating feature between the two viruses being a second RNA segment of unknown function found in MaRNAV-1 and MaRNAV-2 (Charon et al., 2019). Beyond the discovery of these viruses, virtually nothing is known about them.

It can be construed that Matryoshka RNA viruses are specific to the haemosporidian parasite species with which they associate; given the evidence that MaRNAV-1 was only detected in one species of *Plasmodium* (Charon et al., 2019). The discovery of MaRNAV-2, −3, and −4 were performed without the aid of known parasite identification techniques, such as thin-film microscopy and PCR amplification (Bensch et al., 2000; Hellgren et al., 2004). Because of this, the only information available regarding the hosts of these viruses are the definitive avian host species, parasite genus, and parasite lineages. MaRNAV-3, associated with *Leucocytozoon* parasites, was found in only one bird transcriptome (*Acanthis flammea*) co-infected 2 cytochrome *b* lineages of the parasite genus *Leucocytozoon* (Rodrigues et al., 2022). MaRNAV-4, associated with *Haemoproteus* parasites, was detected in two birds of the same species (*Vireo plumbeus*) found to be infected with parasites of the lineages h-VIRPLU01, h-VIRPLU04, and h-TROAE12 (Rodrigues et al., 2022). Because of the multiple lineage infections in these birds, it is impossible to determine the specific parasite-virus association, however, because these viruses have been solely detected in parasite-infected samples, results suggest some specificity to the parasite genus, or perhaps the insect vector.

The prevalence of MaRNAV in current wild avian populations, and their specificity to haemosporidian infection remains unknown. To address this gap in knowledge, we investigated MaRNAV prevalence and diversity using next-generation sequencing and molecular techniques on blood samples collected from mist-net captured California birds, as well as wild birds admitted to a local rehabilitation center. Given that MaRNAV have been consistently detected in association with haemosporidian parasite infection, and the wide geographic distribution of previously detected MaRNAV, we hypothesize that (1) the prevalence of MaRNAV is positively associated with the prevalence of haemosporidian parasites, and (2) MaRNAV infection will be associated with the species of haemosporidian parasite it infects or, alternatively, the insect vector that transmits the parasite. Moreover, we also expected to find novel MaRNAV in this population of birds. Understanding the role of MaRNAV in the avian-haemosporidian parasite system can offer crucial knowledge for future studies on disease ecology and potential implications for bird conservation. Additionally, this research can contribute to a broader understanding of how viruses interact with parasitic protozoans and influence their impact on both avian and human health.

## 2. Materials and methods

### 2.1 Sample Collection

Field samples were collected from four regional parks and two urban parks around the San Francisco Bay Area, CA from October 7, 2022, until April 19, 2024. The locations chosen for sampling were Lake Merced Park, San Francisco, (−122.486302, 37.7130597), Chain of Lakes Meadows, San Francisco, CA (−122.4983867, 37.7660642), Sunol Regional Wilderness, Sunol, (−121.8817683, 37.5200063), Sibley Volcanic Regional Park, Oakland, (−122.2020921, 37.8596852), Tilden Regional Park, Orinda, (−122.2493329, 37.9006318), and Anthony Chabot Regional Park, Castro Valley (−122.0818417, 37.8596852) (Supplementary figure 1). Mist nets were used to capture passerine birds in their natural environment, and 7-10 mist nets were placed at each location. Captured birds were aged and sexed based on morphology (Pyle, 2008), and fitted with an aluminum alloy leg band provided by U.S. Fish and Wildlife. Banding data was submitted to the United States Geological Survey (USGS) Bird-Banding Laboratory. All birds captured were checked for signs of extreme distress, exhaustion, injuries, dehydration, or any abnormalities that would require their immediate release, or need to enter wildlife rehabilitation. None of the field-caught birds appeared to be unhealthy.

A 25-gauge needle was used to extract a blood sample (approximately 50 μL) from the brachial wing vein from each bird. Blood samples were used to make two thin blood smears, and the rest was distributed into two 1.5 mL cryogenic storage tube containing 1 mL of Queen’s Lysis Buffer, for DNA preservation (Longmire *et al*., 1997), and 500 uL of RNA later^TM^ Stabilization Solution for RNA preservation. Samples were kept on dried ice until they could be stored in a - 80°C freezer at the Avian Parasitology Laboratory at San Francisco State University (SFSU). Blood slides were air dried, immersed for around 30 seconds in absolute methanol for fixation, and stained using a 10% Giemsa solution according to the protocols described by Valkiūnas (2005). Slides were examined using a Nikon Eclipse 80i microscope and imaged using QCapture Pro v7.4.4.0 at 100x magnification.

To expand our search beyond passerine birds, additional blood samples were collected from raptors admitted to the Lindsay Wildlife Experience (Walnut Creek, CA) for rehabilitation. Sample collection lasted from November 4, 2022, until May 22, 2024, using the above-mentioned methods to collect thin blood smears, as well as blood stored in Queen’s Lysis Buffer. Samples were collected from the center every 1-2 months and kept at −20 °C, so to account for longer term storage at this temperature, blood was stored in 1 mL of Invitrogen TRIzol LS Reagent^TM^ (Thermo Fisher Scientific, Waltham, MA), instead of RNA later.

### 2.2 Molecular Analysis

DNA was extracted from blood stored in Queen’s Lysis Buffer using the Promega Wizard Genomic DNA Purification System. Purified DNA extracts were stored at −20°C until needed for analysis. Haemosporidian (*Plasmodium*, *Haemoproteus*, or *Leucocytozoon*) presence and identification were detected using a combination of thin film light microscopy and PCR using the primers (HaemNFI/HaemNR3, HaemF/HaemR2, and HaemFL/HaemR2L) and temperatures described by Bensch *et al*. (2000) and Hellgren *et al*. (2004). Amplified PCR products were sent to Elim BioPharm (Hayward, CA) for Sanger sequencing, and sequences were aligned using Geneious Prime® 2024.0.4. Subsequent sequences were cross referenced to GenBank, as well as the MalAvi Avian Haemosporidian Database (Bensch *et al*., 2009)

### 2.3 RNA Seq & Transcriptome Assembly

Blood samples stored in RNA Later were frozen at −80°C until needed for RNA extraction. Before the extraction process, samples were incubated at room temperature for 15 minutes, and centrifuged for 20 seconds at 12,000 x g, and the separated RNA Later was pipetted off. RNA extraction and isolation was performed using the Invitrogen PureLink RNA Mini Kit and treated with on-column PureLink DNase Set. Samples stored in TRIzol were extracted using a phenol-chloroform method, as per the Qiagen RNeasy Mini Kit and treated with Qiagen RNase-Free DNase I. For all samples, a final elution of 30 uL was obtained and stored at −80°C. RNA quality and concentration were assessed using the Agilent 2100 Bioanalyzer at the SFSU Genomics, Transcriptomics, and Analysis Core (GTAC). Samples were chosen for sequencing if they met baseline requirements for RNA Integrity Numbers (RIN) of 5.5 or greater, and concentrations of 20 ng/uL or greater. RNA sequencing was performed by Novogene Co., LTD (Sacramento, CA) using the Illumina NovaSeq PE150 sequencing platform.

A total of 20 samples were selected for sequencing, including 8 *Leucocytozoon*-infected samples, 2 *Haemoproteus*-infected samples, 2 *Plasmodium*-infected samples, and 8 uninfected samples. These samples were chosen at random but ensured that at least one sample representing each of the haemosporidian parasite infection, and a negative control group (uninfected) were included. Paired end reads for each sample were generated in fastq format and released onto a remote server at SFSU. Trimmomatic v0.40 was used to trim adapter sequences from the raw reads, and the Trinity software v2.10.0 was used for *de novo* transcriptome assembly.

### 2.4 Viral Sequence Detection

A homology-based approach was used to detect viral sequences in the assembled transcriptomes, as per Rodrigues *et al*. (2022). Diamond version 0.9.24 BLASTx was used for local sequence alignments against custom BLAST databases consisting of known MaRNAV RdRp protein sequences, as well as a database containing all the known RdRp protein sequences available on NCBI. Hits were then submitted to NCBI BLASTx against the entire non-redundant (nr) database to determine any similar sequences, as well as NCBI BLASTn to determine if the viral elements were integrated into the host’s genomes. Suspected novel RdRp sequences were then submitted to the NCBI ORF finder. The longest ORFs were submitted to the Protein Homology/Analogy Recognition Engine version 2.0 (Phyre2) web portal, as well as HHPred Homolgy Detection Server (Zimmerman *et al*., 2018).

To characterize additional segments of these viruses, a separate Diamond BLASTx database was created that contained MaRNAV-1 and MaRNAV-2 hypothetical proteins discovered by Charon et al. (2019), and the same process as described above was repeated on all transcriptomes from this study, as well as the transcriptomes used by Rodrigues et al. (2022).

### 2.5 Complementary DNA (cDNA) & RT-PCR

Complementary DNA (cDNA) was made from all RNA extracts using the Invitrogen SuperScript IV Reverse Transcriptase (catalog #18090010), using the manufacturer’s protocols. In short, 2 uL of RNA was combined with 1 uL of random hexamers (50 uM; catalog number: N8080127) and 9.8 uL of RNase-free water in a 200 uL RNase-free microcentrifuge tube. This solution was gently centrifuged for approximately 5 seconds, incubated at 65 °C for 5 minutes in a thermocycler, and then on ice for at least 1 minute. A mixture of the following was added to each sample while on ice: 4 uL of the SuperScript IV 5x Reaction Buffer, 1 uL of 10 mM dNTP, 1 uL of 0.1 M DTT, 1 uL of RNaseOUT (40 U/uL), and 1 uL of SuperScript IV Reverse Transcriptase (200 U/uL). This final mixture was gently mixed by pipetting the solution up and down several times and centrifuged for approximately 5 seconds. Samples were then incubated in a thermocycler at 23 °C for 10 minutes, followed by 50 °C for 1 hour, and then 80 °C for 10 minutes. Final cDNA samples were stored at −20 °C until needed for PCR.

To validate the cDNA, oligo primers were created to amplify the phosphoglycerate kinase 1 (*PGK1*) gene, an avian reference gene as described by Olias et al. (2014; Supplementary Data 2), and PCR was performed on all samples to amplify a 450 bp segment of this gene using primers pgk1_F (5’ CACCTTCCTCAAAGTGTCTCA 3’) and pgk1_R (5’ TGAAGTCAACAGGCAGAGTG 3’). The reaction mixture consisted of 25 μL total volume containing 2 μL cDNA template and 23 μL of PCR master mix. The thermocycler profile involved an initial denaturation at 94°C for 5 minutes, followed by 35 cycles of denaturation at 94°C for 30 seconds, annealing at 55°C for 30 seconds, extension at 72°C for 45 seconds, and a final extension at 72°C for 10 minutes.

MaRNAV primers were used as per Charon et al., (2019), and primers were developed for MaRNAV-3, −4, and any novel MaRNAV sequences found via transcriptomics using the Primer 3 Plus Web Interface (Untergasser et al., 2007). All PCR reagents, amounts, concentrations, and temperature profiles are described in supplementary data 1. All amplified products were Sanger sequenced through ELIM BioPharm, as described above, aligned in Genious Prime, and submitted to NCBI BLASTn.

### 2.6 Phylogenetics

Phylogenetics was used to further analyze novel MaRNAV RdRp sequences, and their relationship to all previously described MaRANV, as well as the 20 closest related narnavirus RdRp sequences. Protein sequences were retrieved from NCBI using accession numbers and made into a single fasta file, in addition to the MaRNAV-5 and −6 sequences. The sequences were then aligned using MAFFT v7.309 E-INS-I algorithm, formatted into Phylip|Phylip4, and input into IQ-TREE with the bootstrap value set at 200 (Nguyen et al., 2015). A tree file in Newick format was obtained and loaded to R, and several packages were used to view the tree (ggtree, ggplot2, treeio).

## 3. Results

### 3.1 Sample Collection & Parasite Prevalence

A total of 340 birds were caught using mist-nets in the San Francisco Bay Area (Supplementary Table 1). From these 340 caught birds, 308 blood samples were collected. Additionally, 101 blood samples were collected from the Lindsay Wildlife Museum. This resulted in a total of 409 (n = 409) blood samples used for this study. The overall prevalence of haemosporidian parasite infection in songbirds (field-collected samples) was determined to be 31.17% (n=96) of all mist-net-caught passerine samples, and 23.53% of total (passerine and raptors) blood samples tested. 59 birds were infected with *Haemoproteus,* accounting for 19.16% of passerine birds and 14.23% of total samples, 43 (13.96%, 10.51%) were infected with *Leucocytozoon*, and 19 (6.17%, 4.65%) were infected with *Plasmodium*.

71.29% (n=72) of all raptors were infected with haemosporidian parasites, accounting for 17.60% of total blood samples. This included 62 birds that were infected with *Leucocytozoon* and 12 birds that were infected with *Haemoproteus*. None of the raptor samples were infected with *Plasmodium*.

### 3.2 Novel MaRNAV Detection

Using Diamond BLASTx, there were several hits detected in the transcriptome of an adult male California quail (*Callipepla californica*; accession SAMN43486645) infected with *Haemoproteus lophortyx* (lineage h-COLVIR03; table 1). The top hit had a 71.4% amino acid identity to the RdRp of MaRNAV-4 (E-value = 0.0), and around 40% - 47% identity to MaRNAV-1, −2, and −3 RdRps. The hits varied in length but had high similarity to each other (95% - 100%). The longest transcript sequence was submitted to the NCBI ORF finder, and the longest ORF (2925 nucleotides) was then submitted to Phyre2 and HHpred for a homology-based search. Phyre2 reported the sequence had a 19% identity to an RNA-dependent RNA polymerase with 91.4% confidence, and the HHpred search determined the sequence was an RNA-directed RNA polymerase with 100% probability (E-value = 2.8e-51). These results represent significant evidence of a novel Matryoshka RNA virus, and this sequence was named Matryoshka RNA virus 5 (MaRNAV-5).

**Table 1.** Potential RdRp Sequence Homology Summary.

| Matryoshka RNA Virus | MaRNAV-5 | MaRNAV-6 |
| --- | --- | --- |
| Avian Host | California Quail (*Callipepla californica*) | Barn Owl (*Tyto alba*) |
| Parasite (Lineage) | *Haemoproteus lophortyx* (hCOLVIR03) | *Leucocytozoon californicus* (IBNOW04) |
| Diamond BLASTX Hits & Percent Identity | 71.3% MaRNAV-4 (DAZ89879.1),<br>46.2% Wilkie narna-like Virus 1 (YP_009388589.1),<br>41.7% MaRNAV-3 (DAZ89878.1), | 72.9% MaRNAV-3 (DAZ89878.1),<br>60.4% MaRNAV-1 (QGV56800.1), |
|  | 40.4% MaRNAV-1 (QGV56801.1),<br>44.6% MaRNAV-2 (QGV56804.1) | 62.7% MaRNAV-2 (QGV56804.1),<br>44.4% MaRNAV-4 (DAZ89879.1),<br>43.7% Wilkie narna-like Virus 1 (YP_009388589.1) |
| Nucleotide Length | 2925 nt | 2112 nt |
| Longest ORF | 974 aa | 703 aa |
| Phyre2 Results | 19% Identity to RNA-dependent RNA polymerase (91.4% Confidence) | 33% Identity to RNA-dependent RNA polymerase (88.4% Confidence) |
| HHpred Results | RNA-directed RNA Polymerase (100% Probability, E-value=2.8e-51) | RNA-directed RNA polymerase (100% Probability, E-value=2.2e-31) |

The second positive result was detected in 3 Barn owls (*Tyto alba;* accessions SAMN43486652, SAMN43486653, SAMN43486655) transcriptomes infected with *Leucocytozoon californicus* (l-BNOW04; table 1) (Walther et al., 2016). The top hits shared a 72.9% amino acid identity to MaRNAV-3 RdRp, also detected in a *Leucocytozoon*-infected bird, and 43% - 63% identity to MaRNAV-1, −2, and −4. Similarly, this sequence was submitted to the NCBI ORF finder, Phyre2, and HHpred, and the longest ORF (2112 nucleotides) was determined to be 33% identical to known RNA-dependent RNA-polymerases (phyre2: 88.4% confidence, HHpred: 100% probability, E-value = 2.2e-31). This sequence was named Matryoshka RNA Virus 6 (MaRNAV-6).

Both MaRNAV-5 and −6 were added to the initial reference BLASTx database, and the transcriptomes were re-searched. The MaRNAV-6 RdRp was detected in two other Barn owls that were also infected with *L. californicus*. MaRNAV-1, −2, −3, −4, and −5 were not detected in any of the other transcriptomes.

Using a database of MaRNAV-1 and −2 hypothetical proteins associated with the putative second segment of the virus, similar sequences were discovered in the transcriptome from Rodrigues et al. (2022) in which MaRNAV-3 was discovered, and in all transcriptomes from the current study where MaRNAV-6 were discovered. These sequences were not found in any transcriptome where MaRNAV-4 or MaRNAV-5 were found. The MaRNAV-3 hypothetical protein is 298 amino acids long and has a 29.8% pairwise identity to MaRNAV-2 hypothetical protein (QGV56802.1). MaRNAV-6 hypothetical protein was found to be 302 amino acids long and has a 34.1% identity to MaRNAV-2 hypothetical protein.

### 3.3 Matryoshka RNA Virus Prevalence

All 409 samples were tested for every known MaRNAV RdRp with original oligo primers, as well as primers previously developed by Charon et al. (2019; Supplementary figure 2). 43 of the samples collected from the field were found to be infected with MaRNAV-5, accounting for 44.79% of field samples infected with haemosporidian parasites, 13.96% of all field samples, and 10.51% of all samples used in this study (table 2). All 43 samples were birds infected with *Haemoproteus* parasites, with 13 harboring a co-infection of *Leucocytozoon*, and 1 being co-infected with *Plasmodium*. The virus was detected across 15 different bird species, harboring different *Haemoproteus* lineages that are specific to their definitive avian host. Because of the outliers of birds being co-infected with multiple parasites, we could not make strong association between *Haemoproteus* and MaRNAV-5, despite MaRNAV-5 being found in all *Haemoproteus-* infected birds (χ2 p=0.147).

**Table 2:**
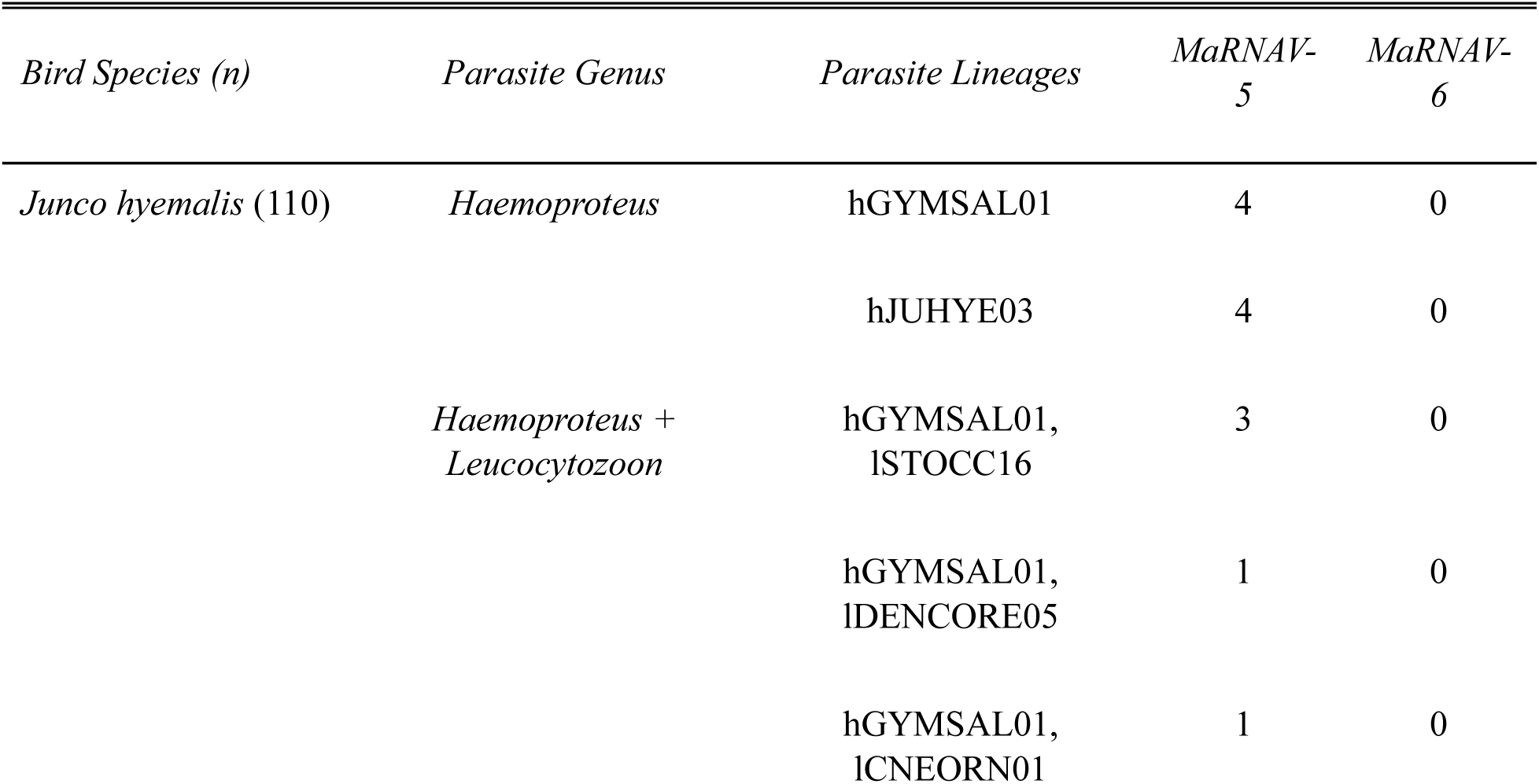
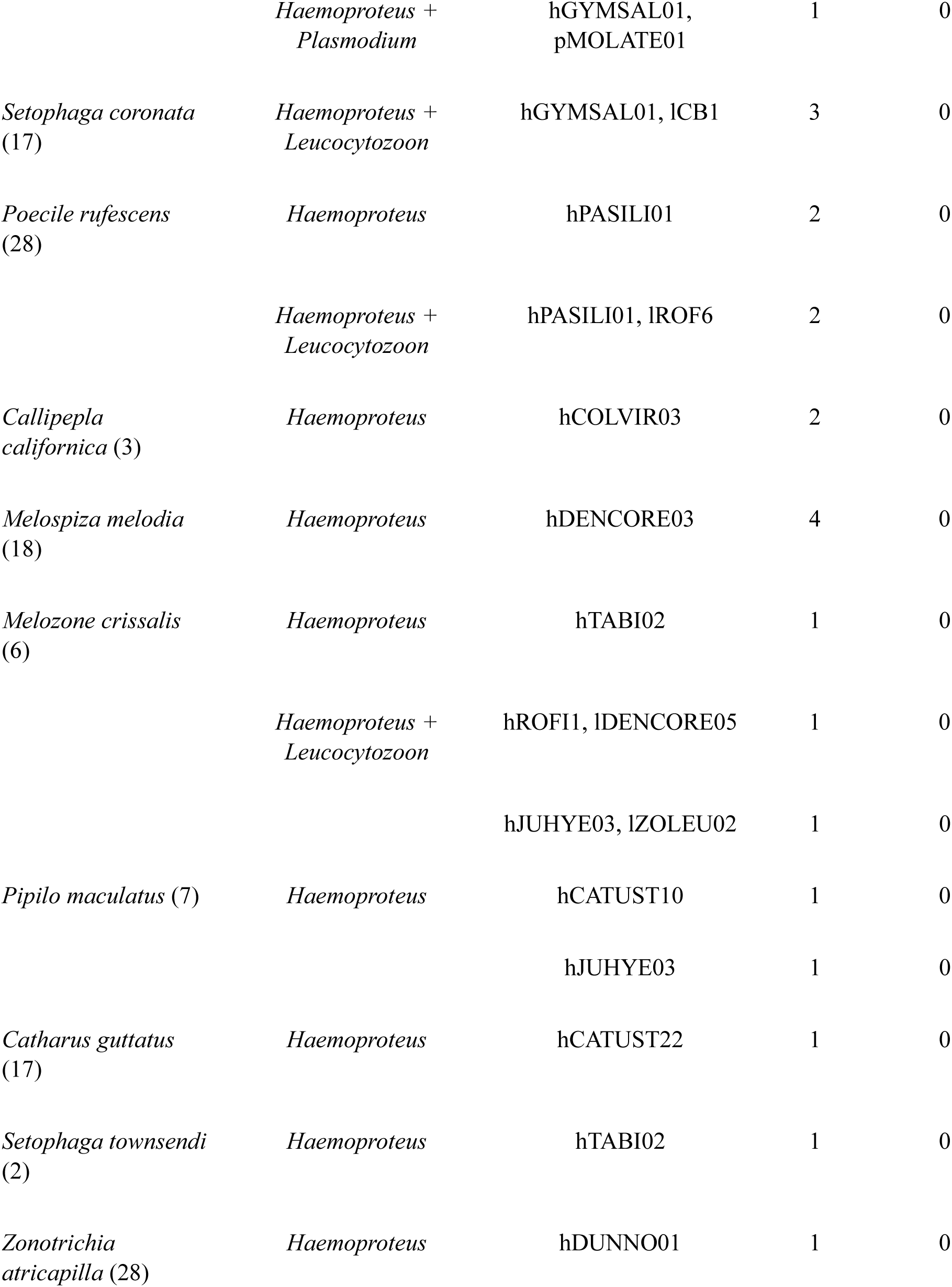
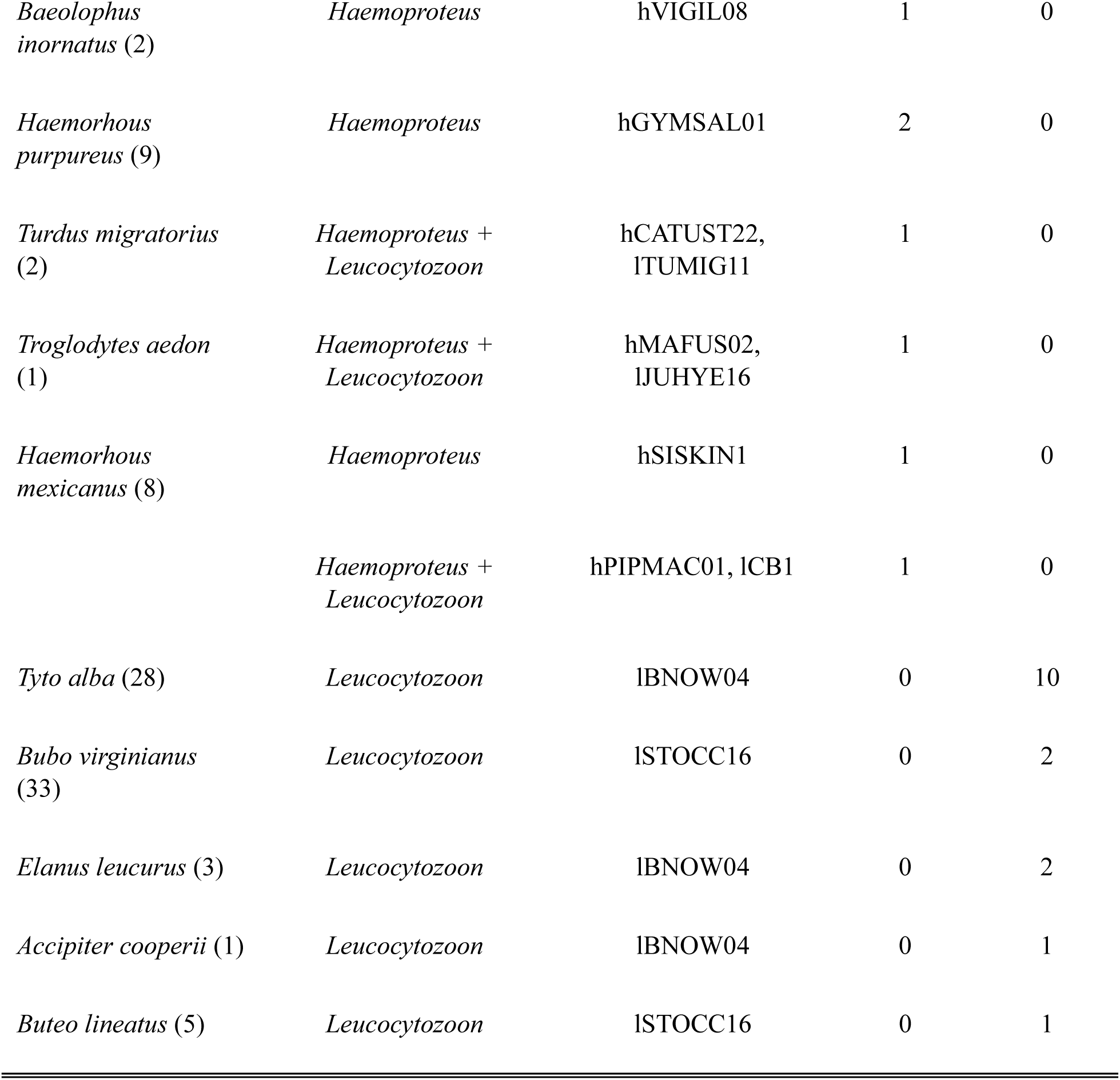
Summary of bird species harboring haemosporidian parasites, their lineages, and MaRNAV-5 or MaRNAV-6.

16 raptor samples were found to be infected with MaRNAV-6, accounting for 22.22% of infected raptors, 15.84% of all raptors, and 3.91% of total samples (table 2). All samples were from raptors that were infected with *Leucocytozoon*, including 2 that were co-infected with *Haemoproteus*. Similarly to MaRNAV-5, MaRNAV-6 was detected across various bird species harboring different *Leucocytozoon*-infections. We found strong association between *Leucocytozoon* and MaNRAV-6 (χ ^2^ p=0.00021).

None of the raptor samples tested positive for MaRNAV-5, and none of the passerine field samples tested positive for MaRNAV-6. No sample tested positive for MaRNAV-1, −2, −3, or −4. Further, MaRNAV presence was seolely detected in haemosporidian-infected bird samples, and no uninfected samples (n=241) ever tested positive for MaRNAV infection, indicating strong association between MaRNAV and haemosporidia infection (χ^2^ p=0). Each sample was tested at least 3 times to test for false positives using cDNA technical replicates made from the same RNA isolates. Only samples that were continuously tested positive for MaRNAV were considered true positives.

### 3.4 Sanger Sequence Analysis

All amplified Sanger sequences showed high nucleotide alignment similarity to MaRNAV-5 and MaRNAV-6 RdRp sequences (95% - 100%), despite being detected across different bird and parasite species (figure 2). MaRNAV-5 was found in birds of different orders, and this assemblage of bird samples also harbored very distinct lineages of *Parahaemoproteus* (table 2). NCBI BLASTx and BLASTn results consistently matched the sequences of MaRNAV-6 and 5 to MaRNAV-3 and MaRNAV-4, respectively.

**Figure 1:**
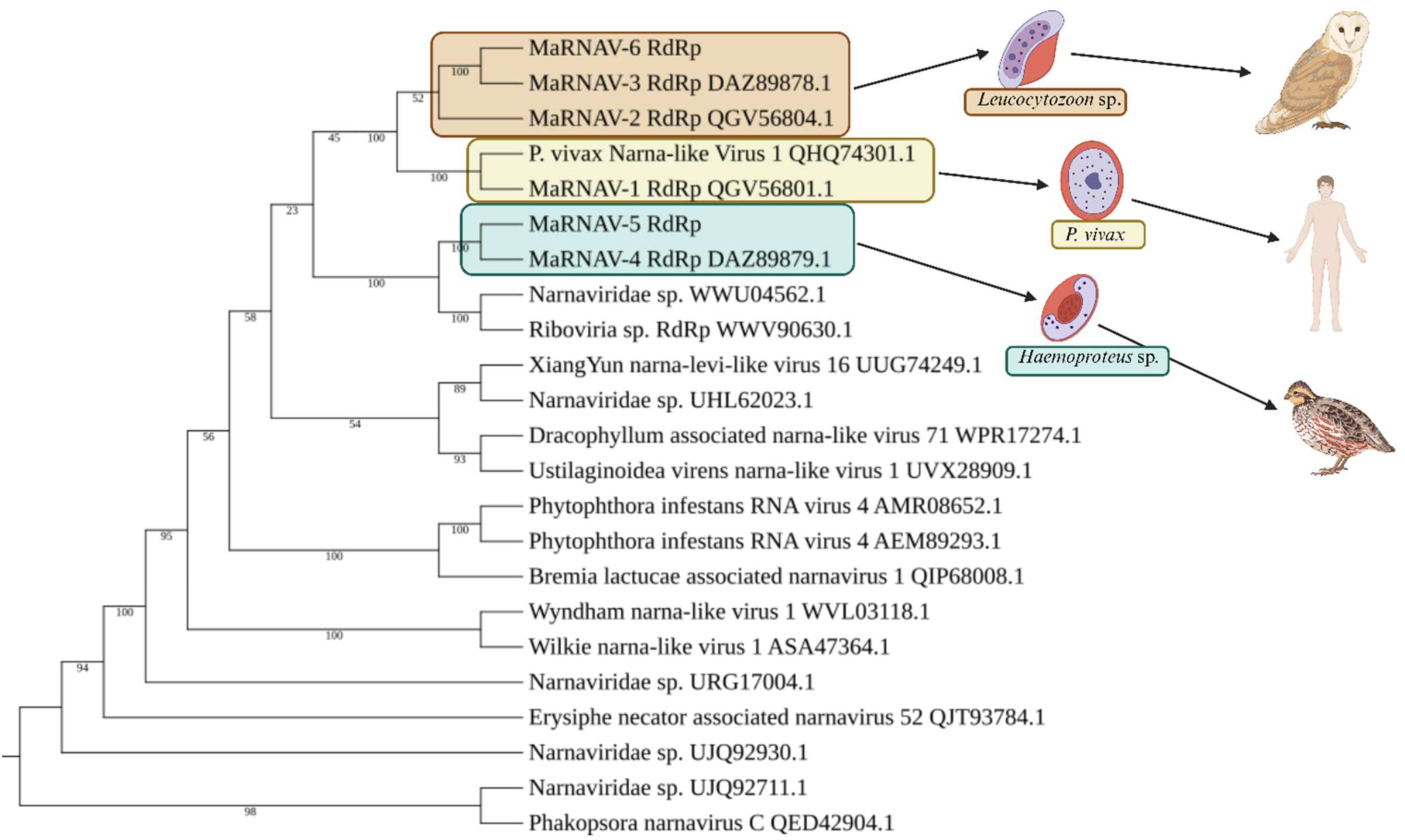
Phylogenetic tree showing how the Protein sequences were aligned using the E-INS-I algorithm in the MAFFT multiple sequence alignment program (v7.309) and put into IQ-tree (v1.6.10) with 200 bootstraps (B=200). IQ-tree chose the LG+F+I+G4 model according to BIC. Edited using BioRender.

**Figure 2:**
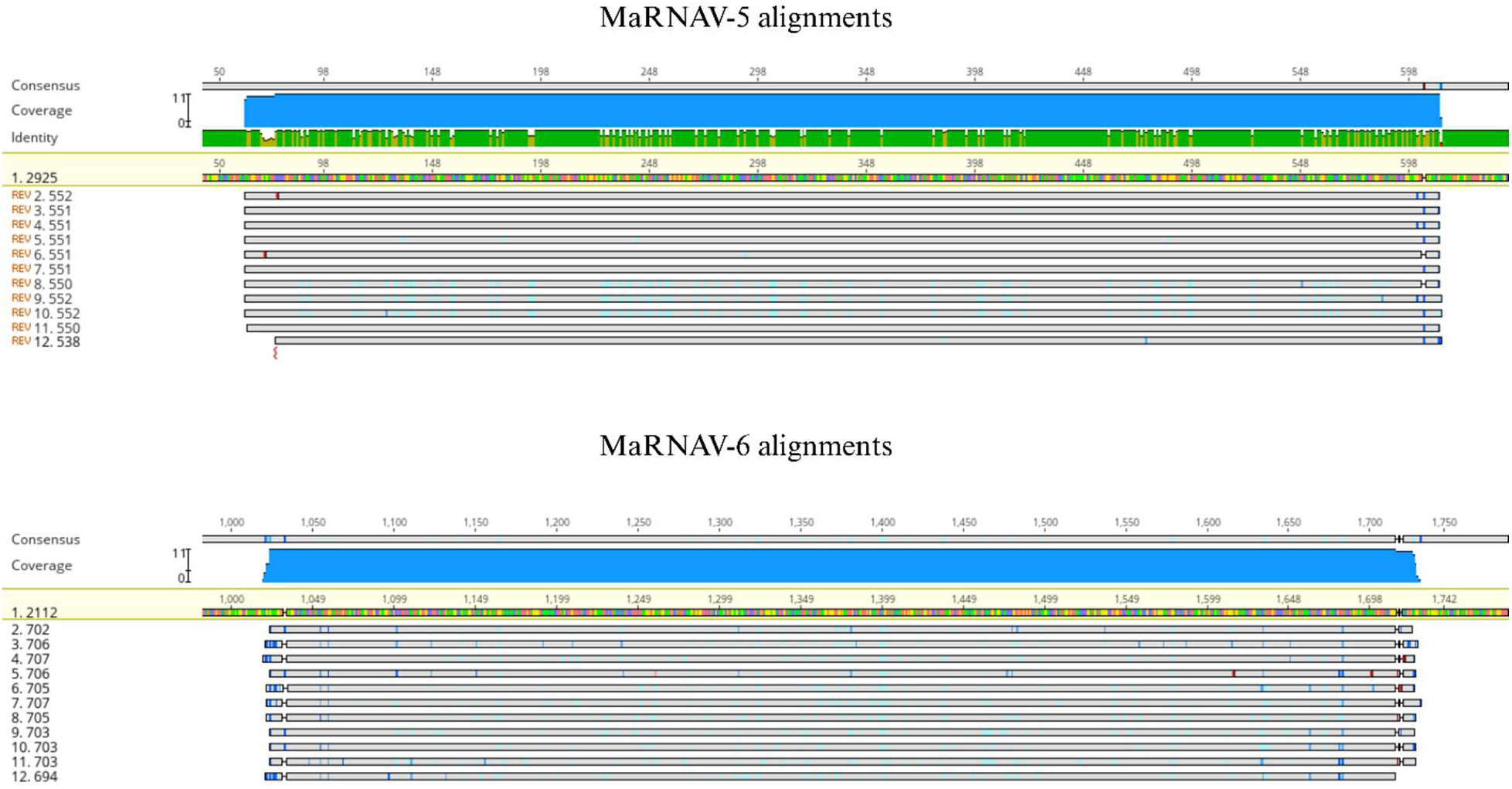
Sanger sequence alignment of amplified PCR products with MaRNAV-5 (top) and MaRNAV-6 (bottom) RdRp sequences. The beige colored lines, labeled as number 1 in both top and bottom, represent the section of the respective RdRp’s amplified by PCR. Each gray line below the RdRp sequence represents a trimmed and aligned sequence that was amplified via RT-PCR. The numbers on the left column represent the nucleotide length for each sequence. The target for MaRNAV-5 primers was a 550 bp sequence, and MaRNAV-6 targeted a 700 bp sequence. Each sequence was trimmed and edited for quality, and alignment was performed using Geneious Prime ® 2024.0.4.

## 4. Discussion

The discovery of haemosporidian parasite-associated viruses raises crucial questions about their potential impact on parasite virulence and host health. Here, we aimed to investigate the prevalence and diversity of novel Matryoshka RNA viruses (MaRNAV) in a wild bird population and determine their association with avian haemosporidian infection. To our knowledge, this study is the first of its kind and provides valuable insights into the prevalence and diversity of MaRNAV in a local avian community. Using molecular methods and transcriptomics, we identified two novel MaRNAV: MaRNAV-5, detected in birds infected with *Haemoproteus* parasites, and MaRNAV-6, associated with *Leucocytozoon* infections. These findings reveal a strong correlation between haemosporidian parasite infection and viral infection (χ^2^ p=0). MaRNAV were exclusively detected in parasite-infected birds, suggesting a close association between the two. These viruses are highly prevalent within haemosporidian-infected bird communities, with between 22.22% and 44.79% of parasite-infected birds harboring these viruses, supporting our hypothesis of viral prevalence correlating with parasite infection.

Phylogenetic analysis of the MaRNAV RdRp region, as well as closely related RdRps, shows that MaRNAV-1, −2, −3, and −6 cluster together, forming their own distinct clade (figure 2). Unsurprisingly, the *Leucocytozoon*-associated MaRNAVs clustered together. The divergent nature of these viruses further prompts a potential reclassification of these viruses as a new genus or viral family. However, MaRNAV-4 and −5 clustered with a narnavirus and ribovirus previously detected in bat metagenomes (WWU04562.1, WWV90630.1). This suggests that the *Haemoproteus*-associated MaRNAV may be more closely related to the canonical narnaviruses, even potentially being narnaviruses themselves. Further evidence of this was our inability to detect the putative second protein segments of MaRNAV-4 or MaRNAV-5, a key characteristic of MaRNAV that distinguish them from narnaviruses, leaving them to be more characteristic of the single-segmented narnaviruses.

### Viral Transmission

An important question pertinent to the biology of the MaRNAV regards how they are transmitted. Previous research suggested that the most likely scenarios involve co-transmission of the virus and the parasite through vertical transmission. In the current study, the detection of novel MaRNAV in multiple avian species, each harboring different parasite species, challenges the previous hypotheses regarding MaRNAV transmission. These results suggest that horizontal viral transmission among wild bird populations may be the main transmission route for MaRNAV. Some potential explanations could account for the patterns observed in this study. The simplest potential explanation is vector-mediated horizontal transmission. The vectors of the haemosporidian parasites (e.g. mosquitoes, biting midges, black flies (Valkiūnas, 2005) may also independently carry and transmit MaRNAV to various bird species.

Here, we posit that the insect is the focal host for the virus, and the virus uses the haemosporidian parasite as a “vector” to shuttle itself to the next insect host (figure 3A). The reason for this hypothesis is that nearly the same viral RdRp sequence can be found in a wide array of bird species, even from different orders. For example, MaRNAV-5 was first found in a California quail (*C. californica*) of the order *galliformes*. RT-PCR then detected the RdRp of MaRNAV-5 in a Dark-eyed junco (*J. hyemalis*) and a California towhee (*M. crissalis*), both of the order Passeriformes. These different bird species and orders carry a broad number of unrelated parasite lineages (supplemental figure 3). However, it is known that both biting midges and blackflies routinely feed on many bird species. Thus, it appears that the virus has developed more specificity towards the insect hosts, which are indeed the definitive hosts of the haemosporidian parasites (Valkiūnas, 2005). A thorough investigation of both the midgut and salivary glands of the invertebrate will be necessary to validate this hypothesis. As an alternative hypothesis, the biting midges that carry and transmit *Haemoproteus*, may independently carry MaRNAV-5 and transmit the virus to various birds already infected with different species of *Haemoproteus* (figure 3B). Likewise with blackflies that carry *Leucocytozoon*. The ecological overlap between the vectors and avian hosts (shared habitats, seasonal activity) may facilitate this hypothetical co-infection process.

**Figure 3:**
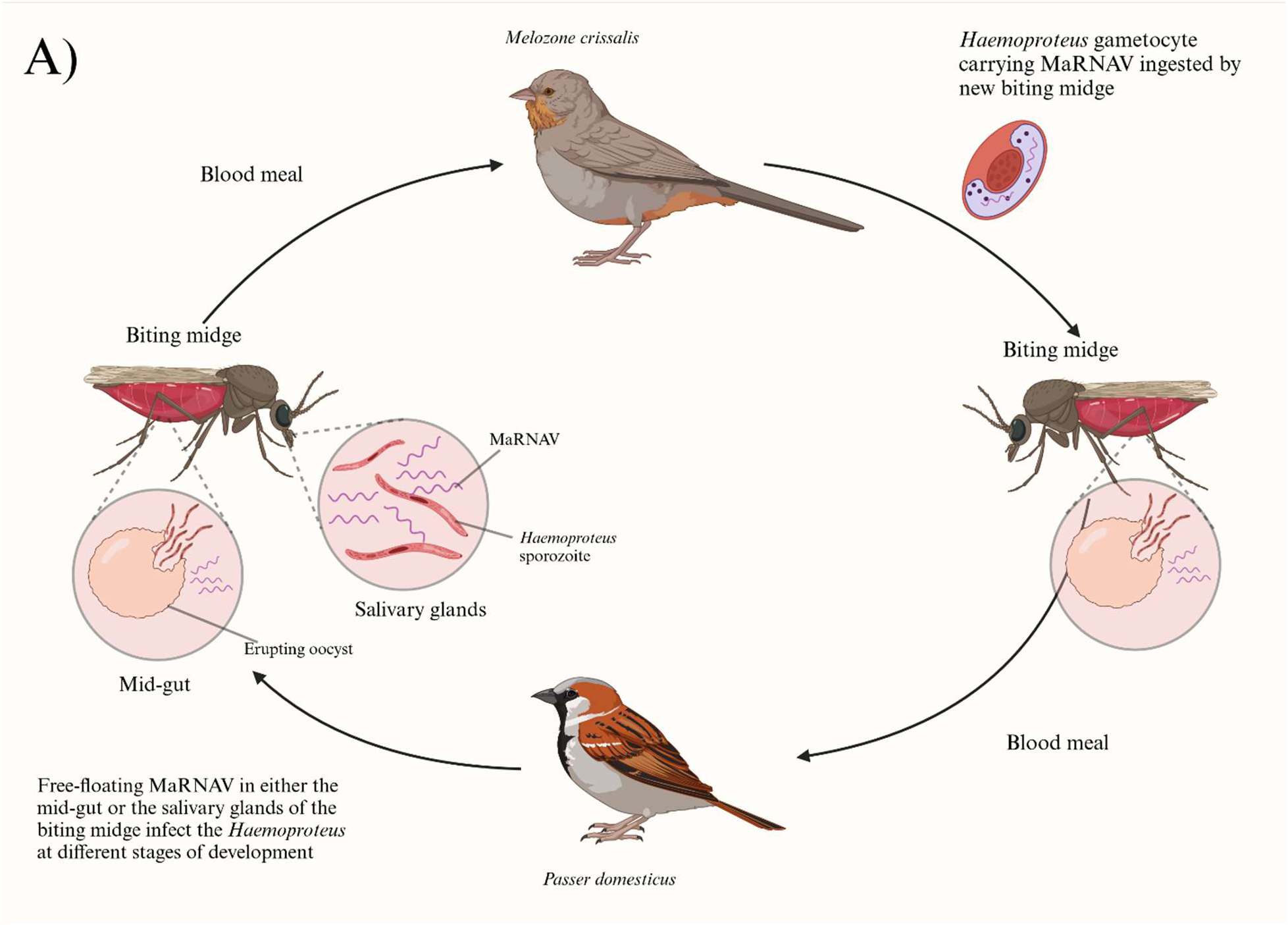
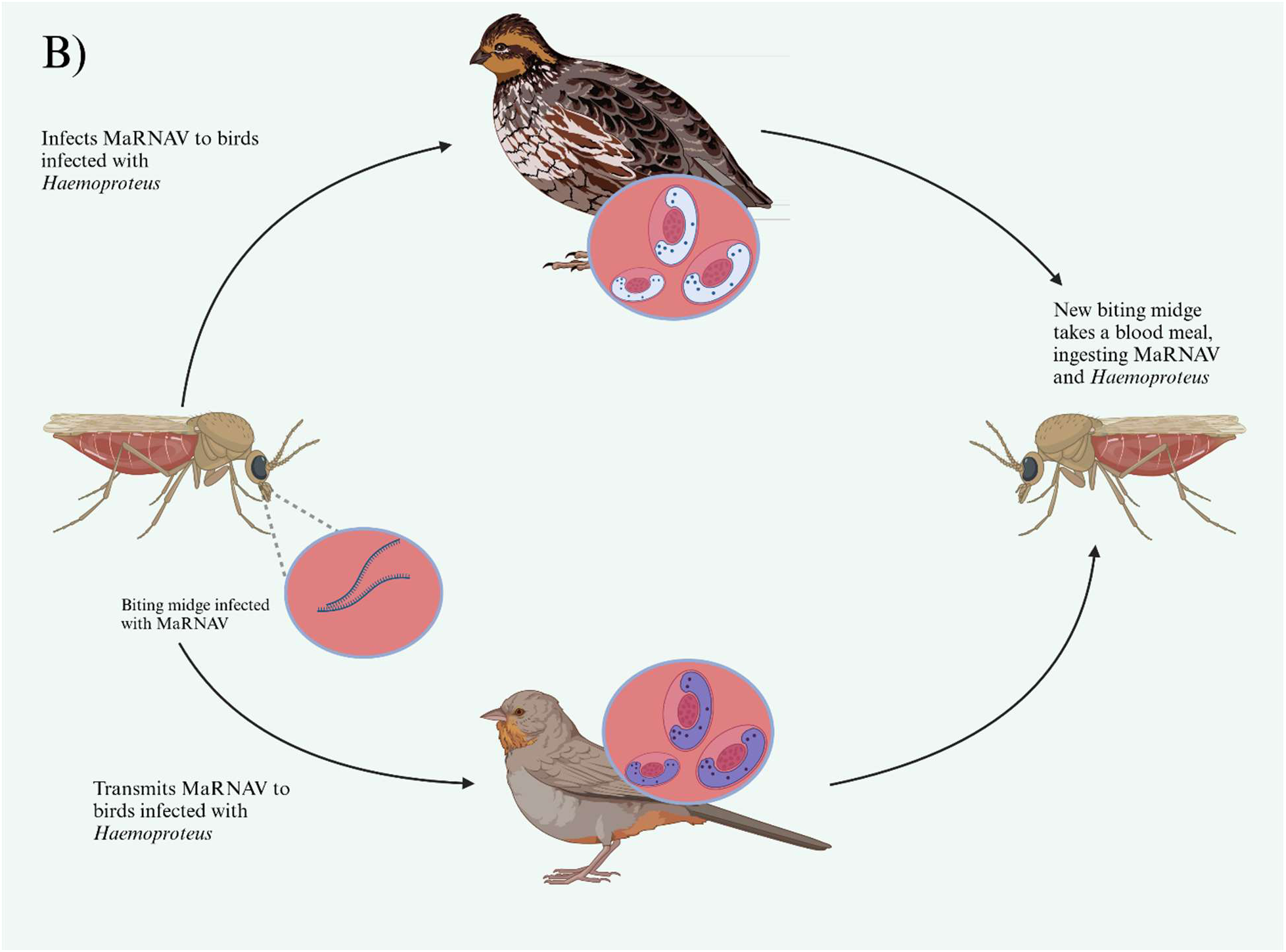
Hypothetical transmission cycle for Matryoshka RNA Viruses, using MaRNAV-5, *Haemoproteus*, and biting midges as an example. A) Depicts the insect vector as the definitive host as the virus and utilizes transmission of the haemosporidian parasites as a “vector” for transmission to the next insect vector. B) Depicts MaRNAV transmission independent of haemosporidian to various avian species infected with various haemosporidian species.

The results from this study, however, point to the first scenario as a much more likely hypothesis. All MaRNAV were found solely in haemosporidian-infected samples, so if an insect vector infected with MaRNAV, were transmitting the virus independent of haemosporidian infection, avian samples uninfected with haemosporidian parasites should have also tested positive for MaRNAV. It is possible, and likely, that this analysis missed other novel MaRNAV, and that the prevalence and diversity of these viruses is much higher than detected in this study. This study was only able to detect novel MaRNAV through meta-transcriptomic analysis, despite searching for previously detected MaRNAV. Given the relatively small sample size (n=20), it is likely that more sequencing would have led to the discovery of more MaRNAV. Genetic diversity generated through recombination or mutation events could also contribute to the MaRNAVs ability to infect a wide range of hosts. This, however, is unlikely given the limited genetic variation found in the RdRp’s detected in this study. If recombination were facilitating viral entry into different hosts, we would expect to find more diverse viral strains.

The effect that these viruses have on their host remains a major unknown. Considering how prevalent they are in nature, it will be essential to study how viral presence affects parasite pathogenicity, if at all. RNA viruses are fast-evolving in nature, narnavirus and narna-like viruses more so due to their simple genomes (Steinhauer & Holland, 1987; Moya et al., 2000; Furio et al., 2005; Duffy et al., 2008). The rapid mutation of RNA viruses can significantly change their functionality, including their virulence, or the parasite’s pathogenicity. In the present study, a single RT-PCR was used to detect the viral presence, suggesting a relatively high titer in the infected blood, while parasite detection typically requires a nested PCR approach. This likely indicates an effect of the virus on the parasite. As it stands, there are essentially three possible means that the MaRNAV can alter the parasite infection in the host: 1) The virus can increase pathogenicity in the host by triggering a type I IFN response, as seen with LRV1, CSpV1, and TVV (Ives et al., 2011; Fichorova et al., 2012; de Carvalho et al., 2019; Rada et al., 2022; Deng et al., 2023). 2) The MaRNAV can decrease the parasite pathogenicity, as with GIV1 (Miller et al., 1998). 3) The MaRNAV does not have any effect on the parasite pathogenicity in the host. These questions may be answered by way of experimental inoculation of birds with MaRNAV and parasite-infected blood and determining gene expression differences between them and healthy birds, and birds infected with haemosporidian parasites without MaRNAV infection. At the surface, the birds captured in this study appeared to be healthy, despite parasite infection.

It is still unknown whether the viruses infect the parasite cell at all or are infecting the animal host cell. All previous data point to the virus being at least associated with haemosporidia, but it has yet to be proven to be infecting the parasite. Uncovering this mystery could potentially give some insight into the transmission and life cycle of the MaRNAV. For example, finding that the MaRNAV does not infect the parasite cells directly increases the possibility of horizontal transmission rather than vertical. Conversely, if the parasite cell is found to be infected with the MaRNAV, vertical transmission is a much more likely transmission mode. Future studies could implement RNA Scope In Situ Hybridization (ISH) to locate and visualize the RdRp within the cell. Recent studies have aimed to visualize *Plasmodium relictum* pSGS1 in avian samples during their dormant parasitemia stages (Himmel et al., 2024). Usage of this technology in the realm of avian haemosporidia opens the possibilities of realistically localizing MaRNAV infections.

The discovery of MaRNAV, and their relationship to haemosporidian parasites could have significant implications for the future of malaria and avian malaria research. If MaRNAV can affect parasite pathogenicity, new strategies may need to be introduced for disease control and prevention. For example, understanding the mechanism in which MaRNAV alter parasite virulence, or host response, can lead to novel therapeutics that target the virus, or the parasite *and* the virus. It is crucial for future work to investigate the insect vector species for the presence of the virus and identify if the virus is more broadly disseminated in the insect’s body. By understanding these complex interactions between MaRNAV, haemosporidian parasites, and their insect vectors, we can gain invaluable insights that may inform future strategies for disease control, prevention, and wildlife conservation.

## 5. Ethical Statement

All field bird handling and sampling protocols and procedures were conducted in accordance with federal, state, and institutional guidelines for the ethical treatment of wildlife. The research was authorized under the United States Department of the Interior, US Geological Survey Permit No. 23555, which allowed for the federal leg banding, taking, possession, and transportation of blood and feather samples, the use of audio lures, baited traps, mist nets, walk-in traps, and other trapping methods. Additional authorization was granted by the State of California Natural Resources Agency Department of Fish and Wildlife under Federal Permit No. S-200730003-20073-001-01. All procedures were reviewed and approved by the Institutional Animal Care and Use Committee (IACUC) at San Francisco State University under Protocol No. A2024-16, titled “Molecular Genetics of Avian Malaria Viruses.” These permits ensured compliance with all relevant ethical standards for the care and use of animals in research, and every effort was made to minimize distress and discomfort to the birds involved in this study.

## 6. Data Availability

Raw RNA sequencing data was submitted to the NCBI SRA under BioProject PRJNA1156234. MaRNAV-5 and -6 sequences have been submitted to GenBank.

## 7. Author Contributions

C.E. performed sample collection, sample processing, data analysis, writing the manuscript text, and prepared all Figures and tables. C.F. Assisted in sample collection, sample processing, and data analysis. S.R. aided in bioinformatic analysis and provided a server in which transcriptomic analysis was performed. RNSM helped develop the project, find funding, provided mentorship, and craft the manuscript.

## 8. Funding

This project was funded by the NIH SFSU/UCSF Bridges to Doctorate Fellowship T32-GM142515, NIH SC3-GM144187 “Molecular Genetics of Avian Malaria Viruses” to RNMS, SC3-GM144187, and S-MIP-23-2 Grant of Research Council of Lithuania to Gediminas Valkiūnas.

## 9. Acknowledgements

We would like to acknowledge the members of the Chaves Lab of Evolutionary Biology at SFSU for their enthusiastic support and help with field collections, especially Jessica Martin and Jaden McCaffrey. We would also like to thank the volunteers and staff at Lindsay Wildlife Experience, located in Walnut Creek, CA, for their tremendous help in obtaining blood samples. A special thank you to Lead Wildlife Rehabilitation Technician, Marcia Metzler RVT, Wildlife Rehabilitation Technicians (in no particular order) Jesse Menne, Aaron Magil, Rachel Komnick, Yani Singer, and Wildlife & Rehabilitation Center Manager Peter Flowers. Thank you to Dr. Gediminas Valkiūnas and Dr. Mélanie Duc from the Nature Research Centre in Vilnius, Lithuania, for their help in haemosporidian species identification, field work, and laboratory assistance. Finally, we want to thank all of the graduate and undergraduate student members of the SFSU Avian Disease Laboratory, student volunteers, as well as the members of the public we met throughout our field work that contributed in any way to helping us. A special thank you to the Avian Disease Laboratory technician Grant Gentile, and Graduate Student Researcher Aileen Lopez.

## 10. Conflict of Interest

The authors declare that the research was conducted in the absence of any commercial or financial relationships that could be construed as a potential conflict of interest.

## 11. Supplementary Material

Supplementary Table 1: Master Sample Collection Metadata

Supplementary Table 2: List of primers used in this study and their function.

**Supplementary Figure 1:**
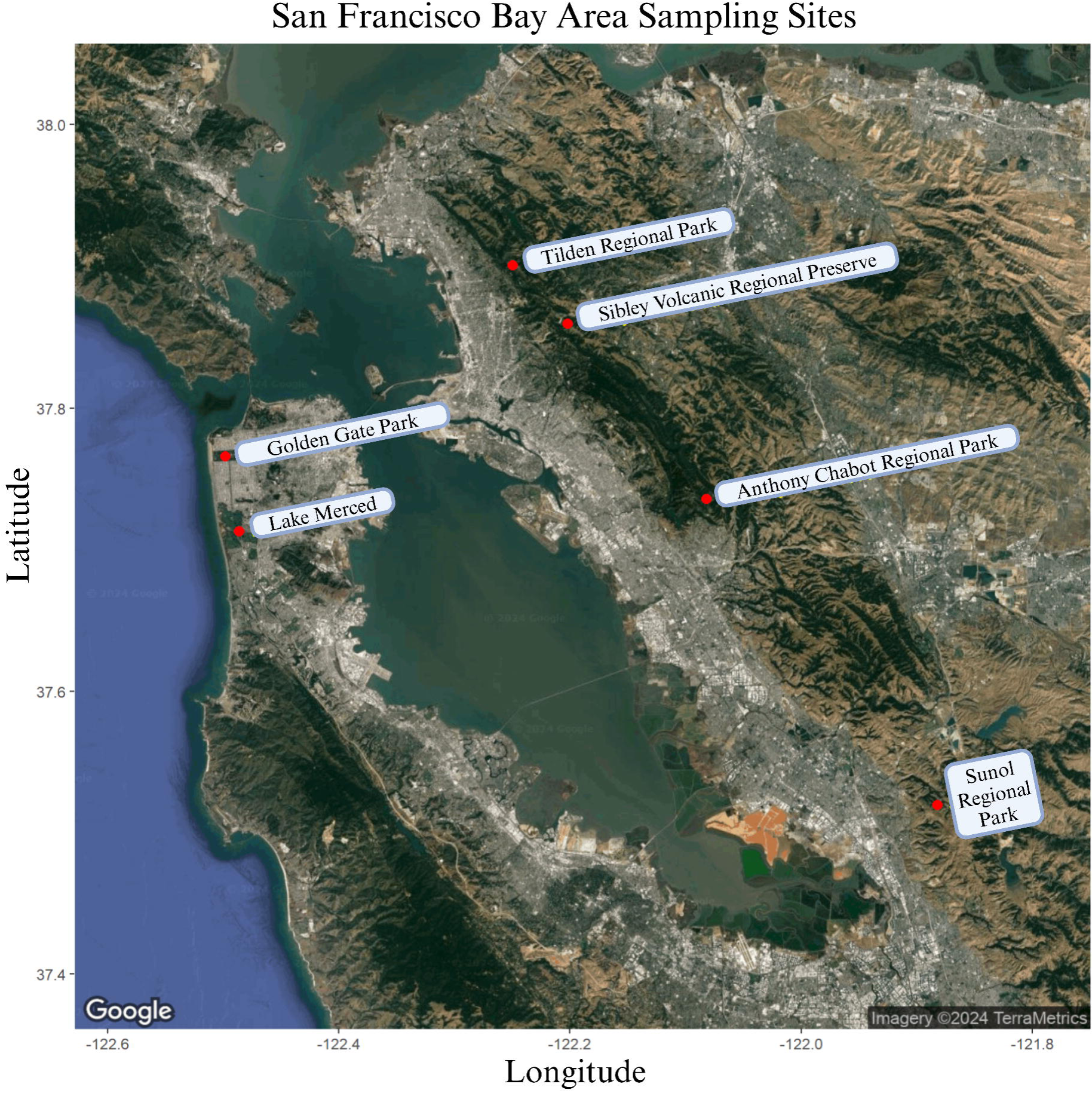
Map of the six sampling sites around the San Francisco Peninsula and East San Francisco Bay Area, CA. The map was created in R (ggmap, ggplot2) using a basemap downloaded from Google maps, and edited using BioRender.

**Supplementary Figure 2:**
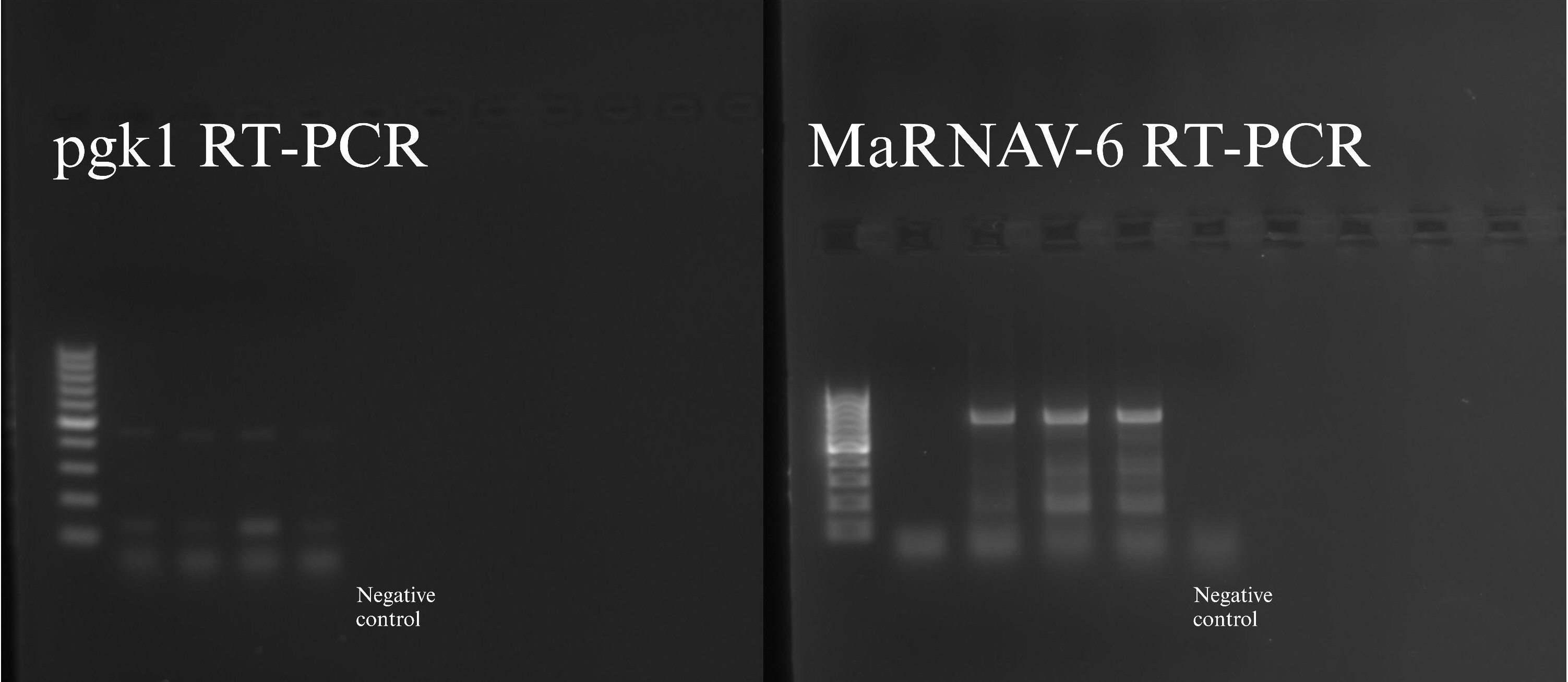
RT-PCR validation of cDNA using primers designed to amplify the *pgk1* gene (left), and MaRNAV-6 RT-PCR results (right). Laboratory grade water was used as a negative control.

**Supplementary Figure 3:**
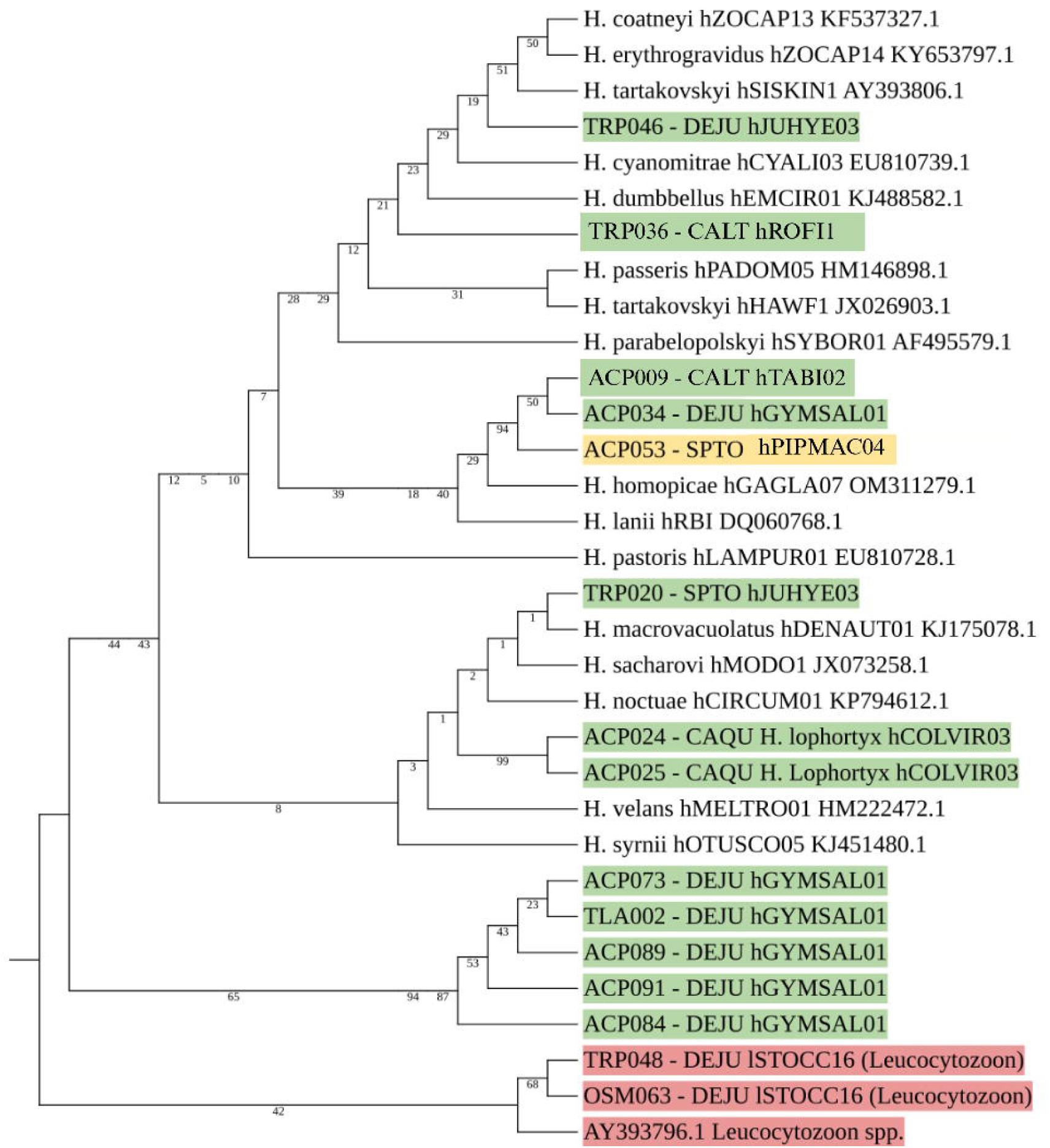
Phylogenetic tree depicting relationship between *Haemoproteus* lineages in this study found to be infected with MaRNAV-5. Labels represent the sample ID, bird host 4-letter ID, and *Haemoproteus* lineage. Green-highlighted labels are samples found in this study to be infected with MaRNAV-5. Yellow-highlighted labels are samples not infected with MaRNAV-5, but novel *Haemoproteus* lineages, submitted to MalAvi and Genbank. Red-highlighted lables represent *Leucocytozoon* outgroup samples. Additional *Haemoproteus* lineages were sequences taken from MalAvi.

## Notes

### Competing Interest Statement

The authors have declared no competing interest.

