## Supplementary material for "Prevalence and Diversity of Haemosporidian-Associated Matryoshka RNA Viruses in a Natural Population of Wild Birds": Suplemental Table 2

| **Supplemental Table 1: Oligo primers used in this study** | | |
| --- | --- | --- |
| Name | Sequence (5’-3’) | Application |
| Pgk1_F | CACCTTCCTCAAAGTGTCTCA | Validation of cDNA |
| pgk1_R | TGAAGTCAACAGGCAGAGTG |  |
| MaRNAV-1_Fw5 | GACTCGTCACCTTGTGAGG | MaRNAV-1 detection (Charon et al., 2019) |
| MaRNAV1_Rev5 | TGGCATCCACTTCAAGCAG |  |
| BW_Narnalike_Fw1 | CTGAAATTGATAARGAYGAAACTC | MaRNAV-2 RdRp (Segment I) detection (Charon et al., 2019) |
| BW_Narnalike_Rev1 | CGTGGCATCCTTYAAATCTGATG |  |
| MaRNAV3_F | AAAGAACAGCCACACCGTTA | MaRNAV-3 RdRp detection |
| MaRNAV3_R | TATACTTCCGCCATGCACAG |  |
| MaRNAV4_F | ATTTATGAGTTCGGGGCCAG | MaRNAV-4 RdRp detection |
| MaRNAV4_R | TGAACCCATGACAAAGCCAT |  |
| MaRANV5_F | TAGGGACGTGTAACCCCAGA | MaRNAV-5 RdRp detection |
| MaRNAV5_R | GAACACTCCCAACCGTGGTA |  |
| MaRNAV6_F | TTTTGGTGGAGCGTGGACATCTT | MaRNAV-6 RdRp detection |
| MaRNAV6_R | GCTCGAGATCCCTGAGTTTC |  |
